# The evolutionary origins of the tumor necrosis factor receptor family – cell death or differentiation?

**DOI:** 10.1101/2019.12.13.875351

**Authors:** Mona Steichele, Angelika Böttger

**Affiliations:** Ludwig-Maximilians-University Munich, Germany

**Keywords:** Hydra, apoptosis, TNF-R superfamily, FADD, ectodysplasin

## Abstract

TNF-R, TNF, and FADD family members are conserved in the fresh water polyp *Hydra.* Moreover, *Hydra* expresses HyTNF-R adaptor proteins similar to the vertebrate TNF-receptor associated factors TRAF-4 and TRAF-6. HyTNF-R is closest related to the human ectodysplasin receptor EDAR, which is involved in epithelial cell differentiation, e.g. the formation of hair and tooth cells in mammals. Consistent with a similar function in *Hydra*, we show here that HyTNF-R protein is localised very specifically in battery cells and in such epithelial cells of the body column that incorporate nematocytes. Epithelial cell differentiation is therefore an evolutionary ancient function of TNF-R/TNF-protein superfamily members. We also show that two *Hydra*-FADD proteins co-localise with *Hydra* caspases possessing death (DD) or death effector (DED) domains in death effector filaments in human cells. Caspase recruitment by members of the FADD-protein family might therefore also be an ancient trait. Future research will have to discover the up-stream pathways, which govern this potential apoptotic pathway in *Hydra* and whether it is extrinsically or intrinsically induced.

## Introduction

Apoptosis describes an essential cellular program for animal development that is crucial for maintaining homeostasis in adult multicellular organisms, to respond to DNA-damage and to promote morphogenesis, reviewed in (Elmore, 2007). The genes governing this program are highly conserved within all animals including cnidarians, nematodes, insects, echinoderms and chordates (Lasi et al., 2010a; Moya et al., 2016).

Apoptosis is regulated by proteases of the caspase family including initiator caspases and executioner caspases, the former being held in an inactive conformation by characteristic pro-domains, e.g. CARDs (**ca**spase **r**ecruitment **d**omains) or DEDs (**d**eath **e**ffector **d**omains). Activation of pro-caspases occurs autoproteolytically by removal of pro-domains and cleavage into small and large subunits, which assemble to form the catalytically active caspase. Activated pro-caspases further activate executioner caspases, which cleave specific substrates to facilitate apoptotic cell death. Pro-caspase activation requires a scaffold (apoptosome), which brings pro-caspases into close proximity (Bao and Shi, 2007). Apoptosomes can be formed in the cytoplasm by interaction of CARD-domains of pro-caspases with APAF-1 (**a**poptotic **p**rotein **a**ctivator 1) in response to intrinsic apoptosis inducers, or at the cell membrane in response to extrinsic apoptotic signals, which are propagated by “death receptors” of the TNF-R (**t**umor **n**ecrosis **f**actor **r**eceptor) superfamily (Salvesen and Riedl, 2008). Binding of TNF-like ligands induces receptor clustering and recruitment of adaptor molecules, e.g. members of the FADD-family (“**F**as **a**ssociated **d**eath **d**omain protein”). FADD-proteins, being composed of DD- and DED-domains cross-link receptor intracellular DDs with DEDs of pro-caspases, increasing the caspase concentration at receptor clusters and allowing auto-proteolysis by induced proximity (Muzio et al., 1998). Mammalian TNF- and TNF-R superfamilies have more than 40 members each, comprising TNF-R-, FasR (CD95), TRAIL-R (**t**umor necrosis factor **r**elated **a**poptosis **i**nducing **l**igand receptor) and EDAR (**e**ctodermal **d**ysplasia **A**-**r**eceptor)-subfamilies for receptors and TNF/FasL-, NGF (nerve growth factor) and EDA-subfamilies for ligands (Wajant, 2003). They function in apoptosis-regulation, inflammation and morphogenesis. In addition to FADDs, adaptor proteins of TRADD- (“**T**NF-**r**eceptor **a**ssociated **d**eath **d**omain protein”) and TRAF- (**T**NF-**r**eceptor **a**ssociated **f**actor) families are recruited to ligand bound TNF-Rs activating NFκB- and TOLL-like or cytoskeletal re-arrangement pathways (reviewed in (Mathew et al., 2009)).

The evolutionary origin of extrinsic apoptotic pathways is unclear. Recent work in the cnidarian *Acropora digitifera* indicated that the TNF-R signalling pathway evolved in pre-cambrian animals. Functional studies exposing human cells to coral TNF and coral cells to human TNF suggested that coral TNF-signalling could be functionally linked to extrinsic apoptosis induction (Quistad et al., 2014). In contrast, genetic approaches in early bilaterian model organisms including *Caenorhabditis elegans* and *Drosophila melanogaster* have so far not revealed any evidence for the existence of a functional extrinisic apoptosis induction pathway (Steller, 2008).

In the cnidarian model organism *Hydra* apoptosis is important for cell number adjustment in response to nutrient supply, oogenesis and spermatogenesis, reviewed in (Böttger and Alexandrova, 2007). Caspases and HyBcl-2 like proteins are involved (Cikala et al., 1999; Miller et al., 2000; Motamedi et al., 2019). The *Hydra* genome also encodes homologs of the mammalian key players of the extrinsic apoptotic pathway, including TNF-R-, FADD- and DED-caspase homologs. In addition, a caspase homolog with a DD-like prodomain (DD-caspase) was described (Lasi et al., 2010a; Lasi et al., 2010b). Here we investigate whether these potential components of a pro-apoptotic signalling module are tied together into a death pathway in *Hydra.*

*Hydra* polyps are composed of two epithelial monolayers, the ectoderm, shielding the organism from the outside and the endoderm, covering the gastric cavity. These are separated by an extracellular matrix, the mesoglea. At the oral end there is a head with tentacles and a mouth opening and at the aboral end there is a peduncle (or foot) terminating in a basal disc. *Hydra* polyps have a high tissue turnover due to constant self-renewal of three stem cell lineages including ectodermal and endodermal epithelial cells and pluripotent interstitial stem cells, residing in interstitial spaces between epithelial cells and giving rise to nerve cells, gland cells and nematocytes. Epithelial cells at the tentacle bases and the basal disc stop dividing and differentiate into tentacle and basal disc cells respectively. Thereby ectodermal tentacle cells enclose nematocytes and become so-called battery cells (Hufnagel et al., 1985). Some epithelial cells in the body column also acquire nematocytes. Battery cells are regularly lost from the tentacles and replaced with new cells arriving from the body column where the epithelium is growing by cell division.

We now show that the single *Hydra* TNF-receptor homolog (HyTNF-R) is very specifically expressed in epithelia cells enclosing nematocytes in all ectodermal epithelia cells in tentacles and in scattered epithelial cells of the body column. Phylogenetically, HyTNF-R is most similar to human EDAR. It interacts with a TRAF-like adaptor, but not with HyFADD-proteins and is probably not involved in extrinsic cell death signalling. HyFADD-proteins, on the other hand, do interact with *Hydra* DED- and DD-pro-caspases.

## Results

### 1. *Hydra* TNF – and TNF-R-homologs are related to human EDA and EDAR

#### Hydra TNF-R (HyTNF-R)

After amplification of the *HyTNF-R*-sequence from *Hydra* cDNA we compared it with sequences of all known human members of the TNF-receptor superfamily (Fig. 1A), which are characterized by two to four extracellular TNF-receptor repeats and an intracellular DD (Aggarwal, 2003). This domain structure was also found in HyTNF-R. With two TNF-receptor repeats HyTNF-R (TNF-R_Hv) was most similar to human EDAR (EDAR_Hs). Phylogenetic analyses of conserved DD-sequences from HyTNF-R and all human TNF-Rs confirmed this (Fig. 1C). More extended phylogenetic analyses using DD-sequences of TNF-Rs from further cnidarians (*Acropora digitifera, Nematostella vectensis), Danio rerio, Xenopus laevis, Xenopus tropicalis* and *Mus musculus* revealed a branch of cnidarian TNF-R DD-sequences that were most closely related to vertebrate EDAR- and NGF-receptor (p75) sequences (Fig. 2). This relationship is supported by previous phylogenetic analyses revealing several *Acropora* TNF-receptor sequences on a branch with human EDAR (Quistad et al., 2014).

**Fig. 1.**
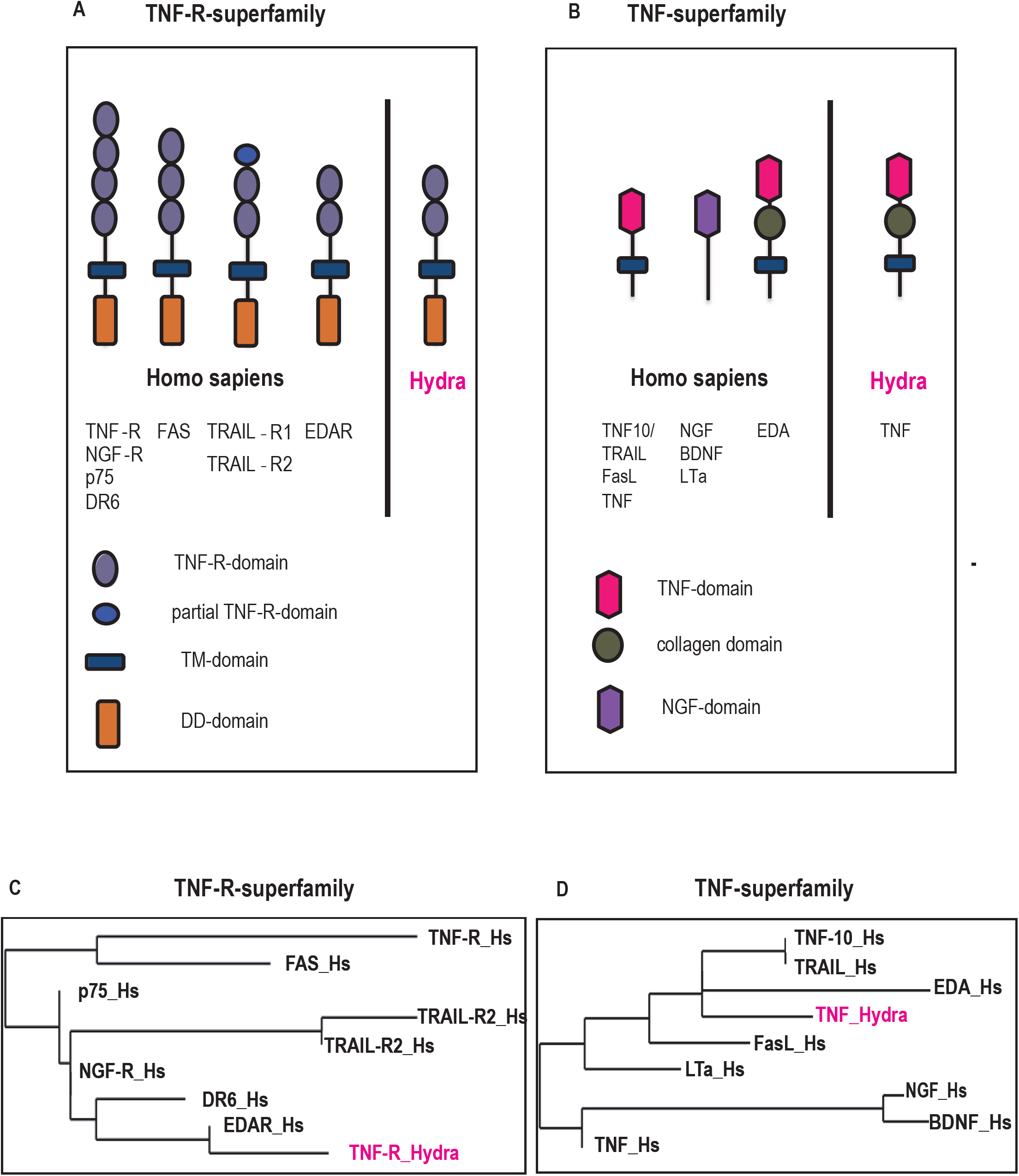
Comparison of Hydra TNF-R and TNF with human homologs: (A): Comparison of the domain structure of human DD-domain containing members of the TNF-R superfamily with HyTNF-R. (B) Comparison of the domain structures of human members of the TNF superfamily and HyTNF. (C) Neighbor joining phylogenetic tree of *Homo sapiens* [Hs] and *Hydra vulgaris* [Hv] TNF-R superfamily members. Only DD-domain sequences were used. TNF-R[Hs]: NP_001333021.1, p75[Hs]: NP_002498.1, NGF-R[Hs]: AAB59544.1, EDAR[Hs]: AAD50077.1, FAS[Hs]: AKB11528.1, TRAIL-R1[Hs]: NP_003835.3, TRAIL-R2[Hs]: AAC51778.1, DR6[Hs]: NP_055267.1, TNF-R[Hv]: XP_004209206.1 (D) Neighbor joining phylogenetic tree of human TNF superfamily members and HyTNF. TNF/NGF domain-sequences were used of human and hydra ligands. TNF[Hs]: NP_665802.1, TNF10[Hs]: NP_003801.1, Eda[Hs]: AAI26144.1, NGF[Hs]: AAH32517.2, FasL[Hs]: AAO43991.1, TRAIL[Hs]: NP_003801.1, LTa[Hs]: XP_011512918.1, BDNF[Hs]: CAA62632.1, TNF[Hv]: XP_012554653.1

**Fig. 2.**
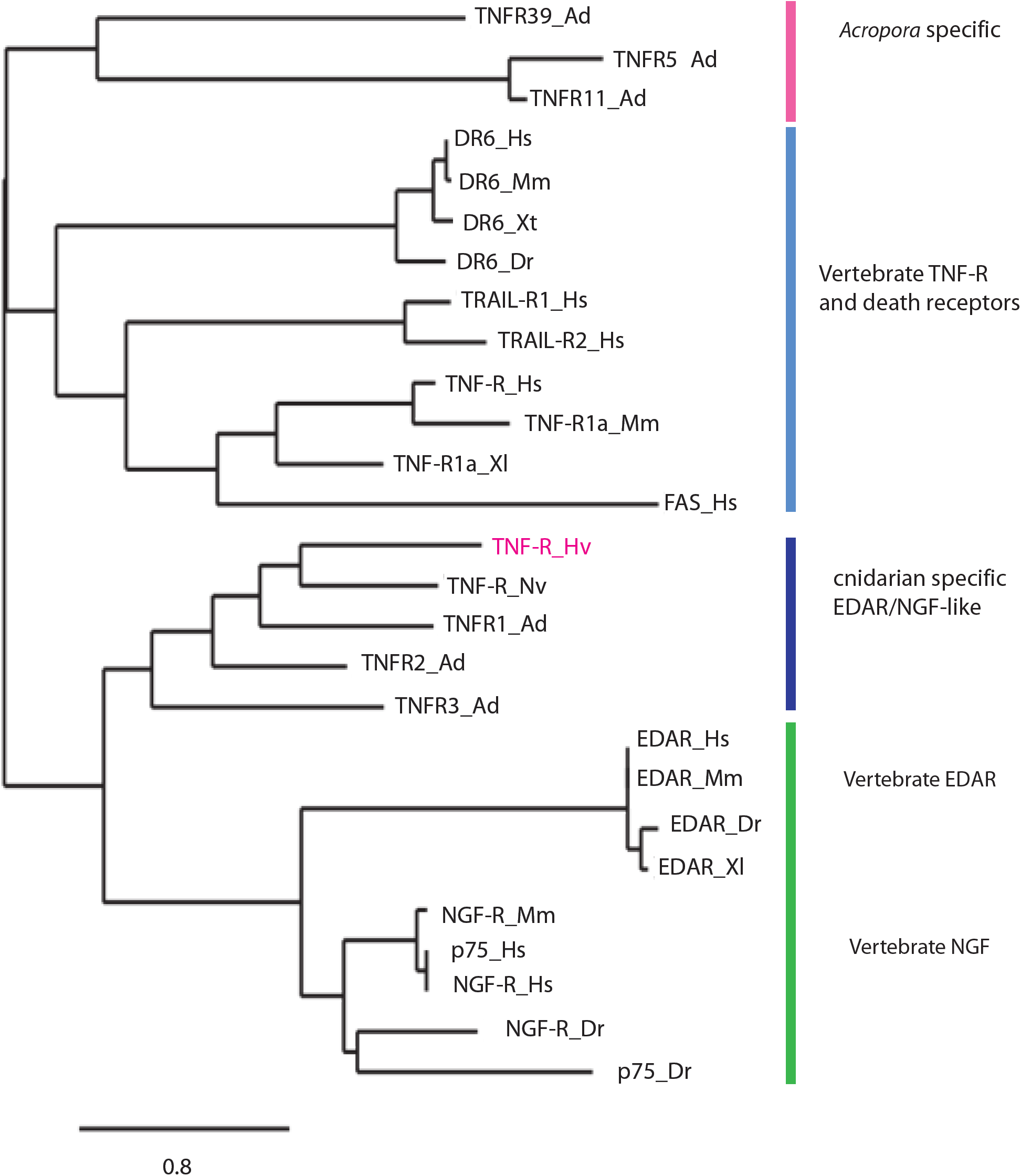
Neighbor joining phylogenetic tree of TNF-R family members: Death domain sequences of the following TNF-R superfamily members were used: *Homo sapiens* [Hs], *Mus musculus* [Mm], *Danio rerio* [Dr], *Xenopus laevis* [Xl], *Acropora digitifera* [Ad], *Nematostella vectensis* [Nv] and *Hydra vulgaris* [Hv]. TNF-R[Hs]: NP_001333021.1, p75[Hs]: NP_002498.1, NGF-R[Hs]: AAB59544.1, EDAR[Hs]: AAD50077.1, FAS[Hs]: AKB11528.1, TRAIL-R1[Hs]: NP_003835.3, TRAIL-R2[Hs]: AAC51778.1, DR6[Hs]: NP_055267.1, DR6[Mm]: AAK74193.1, EDAR[Mm]: XP_006513267.1, TNF-R1a[Mm]: AAH52675.1, NGF-R[Mm]: AAD17943.1, EDAR[Dr]: ABP03881.1, p75[Dr]: XP_700985.3, NGF-R[Dr]: NP_001185589.1, DR6[Dr]: ABG91568.1, EDAR[Xl]: NP_001080516.1, DR6[Xt]: ABQ51095.1, AdTNFR1[Ad]: 12827Acd, AdTNFR2[Ad]: 07010Acd, AdTNFR3[Ad]: 11053Acd, AdTNFR5[Ad]: 14243Acd, AdTNFR11[Ad]: 09194Acd, AdTNFR39[Ad]: 08700Acd, TNF-R[Nv]: XP_001625370.1, TNF-R[Hv]: XP_004209206.1; *Acrospora* non EDAR-like sequences (pink), vertebrate TNF-R superfamily members outside EDARs (light blue), Cnidarian TNF-Rs close to vertebrate EDARs (dark blue) and vertebrate EDAR- and NGF-R-like sequences (green).

#### Hydra TNF (HyTNF)

Human EDAR is activated by EDA, a member of the TNF-superfamily possessing a collagen domain in addition to its extracellular TNF-domain (Mikkola and Thesleff, 2003). By BLAST-searching the *Hydra* genome with the TNF-domain sequence of human TNFα, we found a putative HyTNF encoding a C-terminal TNF-domain, a collagen-like domain and a transmembrane domain. This domain structure was the same as in human EDA, but different from the domain structures of human TNF, FasL or NGF, which all lack a collagen domain (Fig. 1B), (Swee et al., 2009). Phylogenetic analysis of the conserved TNF-domains of all human TNF-related sequences confirmed that HyTNF (TNF_Hv) was most similar to human EDA (Fig. 1D).

#### Hydra TNF-receptor protein is localised in epithelial cells that incorporate nematocytes

In order to investigate the function of HyTNF-R we raised an antibody against a peptide comprising 22 amino acids of the DD of the protein (Davids Biotechnology, Regensburg, Germany). This antibody detected GFP-tagged HyTNF-R introduced into human HEK293T-cells on SDS-PAGE/Western blots (Fig. S1A). It was also able to precipitate HyTNF-R from Hydra cell lysates (Fig. S1B). Moreover, it specifically stained HyTNF-R that was overexpressed in human HEK-cells in immunofluorescence experiments (not shown).

Immunofluorescence on *Hydra* whole mounts revealed staining of scattered ectodermal epithelial cells in the body column and of all battery cells in the tentacles (Fig. 3A-D). A closer look at the epithelial cells in the body column showed that each of them had incorporated one or two nematocytes (Fig. 3D, E). We also stained thinly cut sections of *Hydra* polyps with this antibody to confirm the presence of nematocytes within TNF-R positive ectodermal epithelial and battery cells (Fig. 3 F-I). The middle section of a polyp expressing GFP in all endodermal cells depicts TNF-R-staining in five scattered ectodermal epithelial cells of the body column and in tentacle battery cells. Neamtocytes are clearly recognisable by the outlines of capsule structures and the typically half-moon shaped nuclei of nematocytes harbouring these capsules (see magnifications in Fig. 3G-I).

**Fig. 3.**
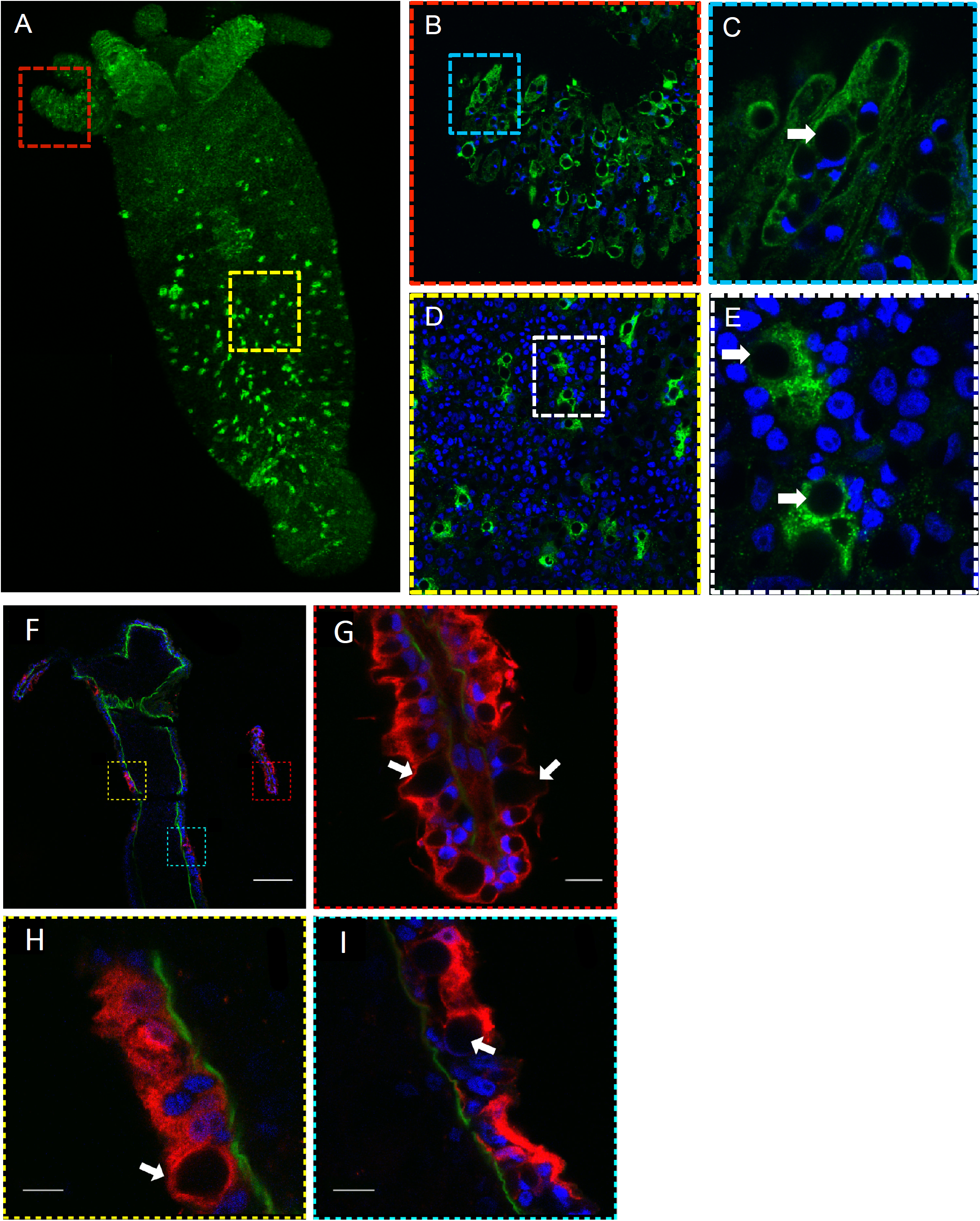
Localisation of HyTNF-R: Laser confocal microscopic images of *Hydra* whole mounts stained with anti-HyTNF-R antibody. (A) Overview whole polyp (B) Tentacle (C) Battery cell (D) Part of the body column (E) Two ectodermal epithelial cells with incorporated nematocytes (white arrows point at stenotele capsules). (F) Thin section of *Hydra* polyp expressing GFP in endodermal cells (animals were a kind gift from Rob Steele, Irvine), GFP-signal (green), anti-TNF-R antibody-signal (red) and DNA-signal (DAPI, blue). Yellow, red and green boxes indicate positions of magnified images in G (tentacle battery cell, red box), H and I (capsule enclosing cells in body column, yellow and green boxes), scale bar 100μm, white arrows point at nematocyte capsules

When we expressed HyTNF-R-GFP in single *Hydra* cells via biolistic transformation (Fig. 5A) GFP-signals were found in vesicular structures in the cytoplasm and at the plasma membrane. We did not find any signs of apoptotic cells.

**Fig. 5.**
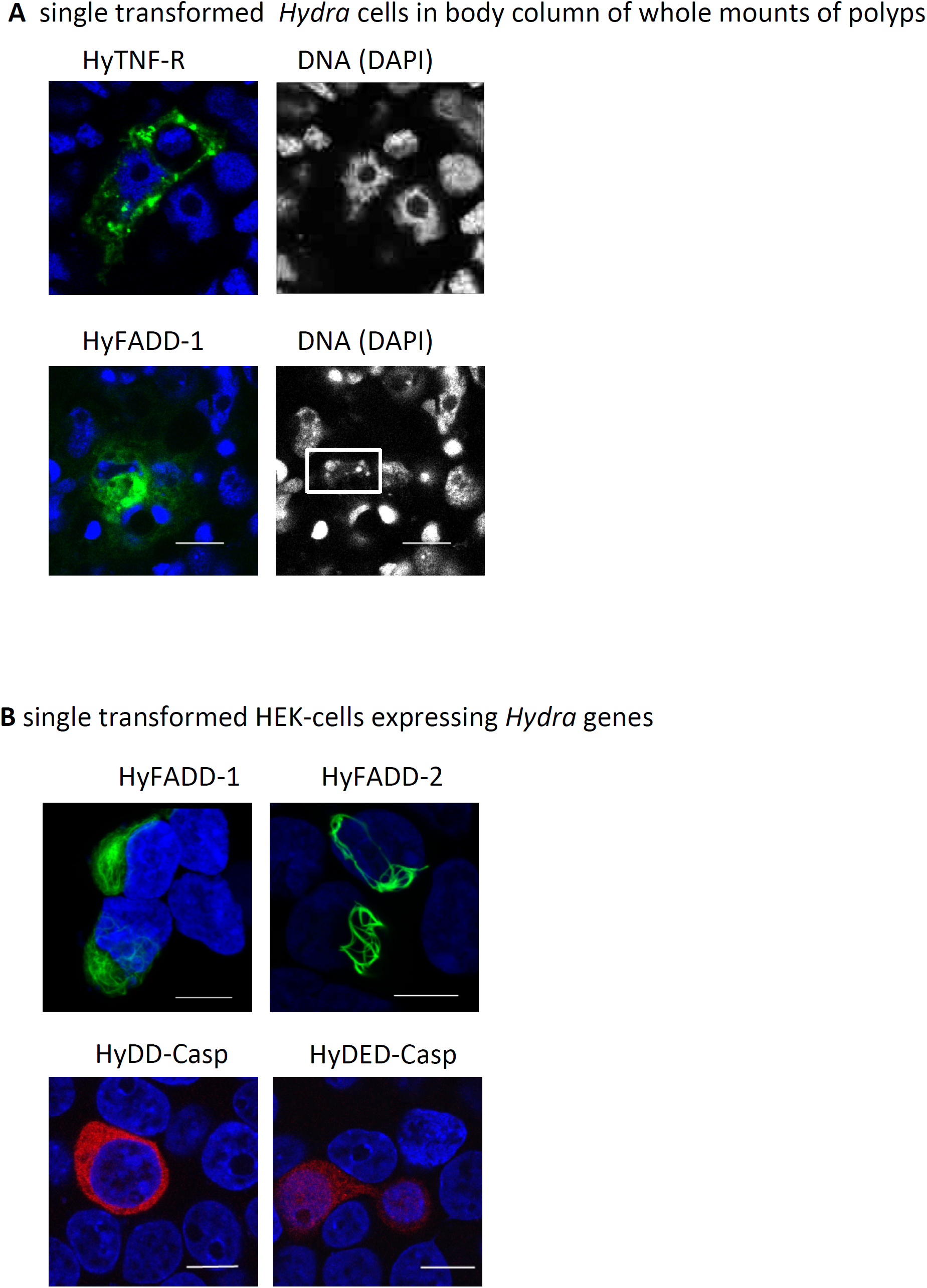
Overexpression of proteins in *Hydra* and in human HEK293T-cells. Laser confocal microscopic images of (A) GFP-tagged HyTNF-R and HyFADD-1 in *Hydra* cells after biolistic transformation as indicated. Left hand panels show merged images of GFP and DAPI signals, right hand panels show DAPI signal in grey. White rectangle: apoptotic nucleus of a *Hydra* cell. (B) GFP-tagged HyFADD-1, HyFADD-2 and HA-tagged HyDD-Casp and HyDED-Casp in HEK293T cells as indicated. Merged images of GFP and DAPI signals or anti-HA-antibody and DAPI signals are shown.: GFP (green) anti-HA antibody (red), DAPI (blue); Scale bars: 10 μm.

#### HyTNF-R interacting proteins

In order to find proteins that interacted with HyTNF-R we used the anti-TNF-R antibody for immunoprecipitation of *Hydra* cell lysates and subjected the precipitate to quantitative mass spectrometric analysis. Table S1 lists 30 proteins that were identified with a probability score of 100% in the samples of proteins co-precipitated with anti-HyTNF-R antibody, but not with the control antibody. Functional assignments indicated co-precipitation of a *Hydra* TRAF-6-like protein, and a homolog of an IκBkinase subunit. Furthermore, more than half of the co-precipitated proteins could potentially interact with HyTNF-R containing protein complexes during its biosynthesis, trafficking and endocytotic recycling. This includes cytoskeletal proteins associated with actin or tubulin dynamics, three membrane proteins, three ECM (extracellular matrix) proteins and a substantial number of proteins involved in translation and protein sorting. HyFADD was not detected (Fig. 4A).

**Fig. 4.**
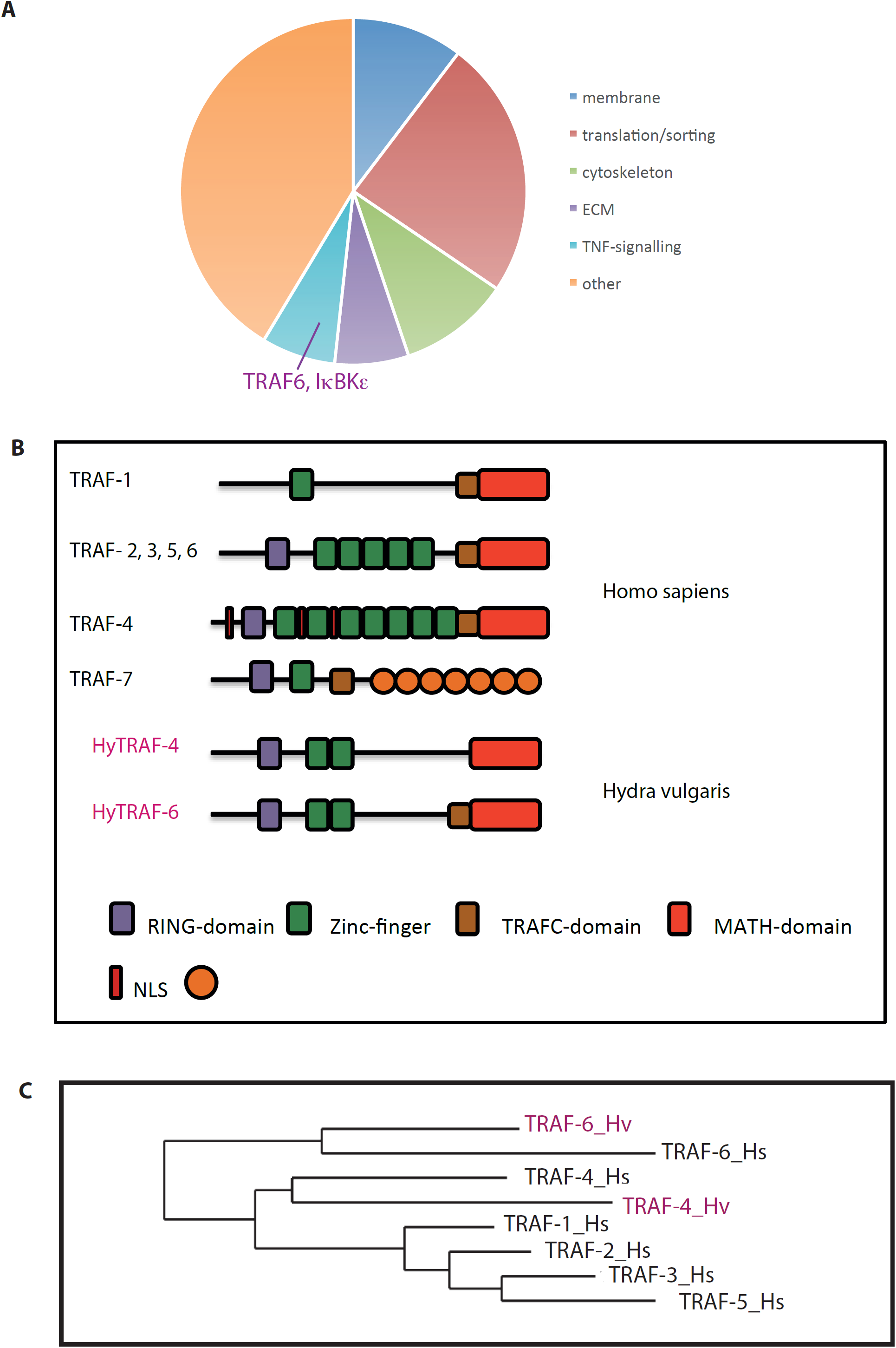
Hydra TRAF-proteins. (A) Diagram indicating functional classification of proteins specifically precipitated with anti-HyTNF-R antibody. Only proteins identified with 100 % probability were included; “others” refers to functional assignments of precipitated proteins that we can not connect to TNF-R at the moment, TRAF-6 and IκBK subunit ε-homologs are indicated in pink letters (B) Representation of domain structures for human and *Hydra* TRAF-proteins; Human TRAFs 2, 3, 5, 6 have the same domains but only one example is given. (C) Neighbor joining phylogenetic tree of *Homo sapiens* [Hs] and *Hydra vulgaris* [Hv] TRAFs based on the sequences of the MATH-domains. TRAF1[Hs]: NP_001177874.1, TRAF2[Hs]: ADQ89802.1, TRAF3[Hs]: NP_663777.1, TRAF4[Hs]: NP_004286.2, TRAF5[Hs]: NP_001029082.1, TRAF6[Hs]: NP_665802.1, TRAF4[Hv]: XP_004207529.1, TRAF6[Hv]: XP_004206538.1

We next investigated the possibility that HyTNF-R uses a *Hydra* TRAF-protein as an adaptor. We therefore cloned the sequences of HyTRAF-6 (T2M357_HYDVU) and HyTRAF-4 (which we identified by additional database search, (XP_004207529.1)) from *Hydra* cDNA. Sequence analyses and protein domain-searches revealed that they have an N-terminal ring finger, two TRAF-type zinc fingers and a MATH-domain (**m**eprin-**a**nd **T**RAF-**h**omology), which is responsible for the interaction of TRAFs with receptors (Fig. 4B). TRAF-6 additionally has a coiled coil region. Comparison of HyTRAF-4 and HyTRAF-6 domain structures with those of human TRAFs revealed similarity of human TRAF-2, 3, 5 and 6 with HyTRAF-6, except that the *Hydra-*protein has only two instead of four Zinc-finger domains. HyTRAF-4 with its lack of the coiled-coil-domain has a domain structure that does not occur in human TRAFs. However, phylogenetic analyses comparing the protein domains and the sequences of the conserved MATH-domains from human and *Hydra* revealed that human TRAF-4 and TRAF-6 are related to HyTRAF4 and HyTRAF6 respectively, whereas all other human TRAF-sequences (TRAF-1, 2, 3, 5) form a separate group indicating that they might have evolved later (Fig. 4C). This is supported by a previous study, which had suggested TRAF-4 and TRAF-6 as the founding members of the family (Chung et al., 2002). The large differences in the domain structures of TRAF-proteins probably reflect variation in signalling cascades to which they can be linked.

In summary these data suggest that *Hydra* TNF and TNF-R are similar to mammalian EDA and EDAR and that HyTNF-R could potentially use HyTRAF-4 and 6 proteins as adaptors. We propose that it has a function for epithelial cell differentiation into battery cells, but not in apoptosis. In order to search for alternative possibilities for an extrinsic apoptotic pathway in *Hydra*, we investigated *Hydra* FADD-proteins.

### 2 A potential HyFADD-DED/DDcaspase-cell death pathway

#### HyFADD1 and HyFADD2 form death effector filaments

We cloned two potential FADD-genes from *Hydra* cDNA and named them HyFADD-1 (XP_002166467.1) and HyFADD-2 (XP_002166848.2). Both HyFADD-protein sequences contained a DD followed by a DED (Lasi et al., 2010a). This domain structure is identical with that of human FADD, but different from the domain structures of human TRADD and EDARADD (not shown). When we biolistically transfected *Hydra* cells with plasmids encoding HyFADD-1 and HyFADD-2 we did not see any epithelial cells expressing these proteins. In the case of HyFADD-1 we pictured two cells with HyFADD-GFP-signals, which were clearly apoptotic (Fig 5A, S2). These experiments suggested that HyFADDs might induce apoptosis in *Hydra* cells. We then expressed GFP-tagged HyFADD-1 and HyFADD-2 in human HEK cells. Here the transfected cells accumulated both HyFADD proteins in long cytoplasmic fibres. For HyFADD-1 these resembled a cage around one half of the nucleus. HyFADD-2 filaments were very elongated throughout the cytoplasm (Fig. 5B). Similar fibres had previously been observed with human FADD-proteins in HeLa-cells. These had been designated DEFs (“death effector filaments”) by Siegel et al. (Siegel et al., 1998).

#### HyDDCasp and HyDEDCasp are recruited to FADD1 and 2 death effector filaments

We then investigated the distribution of HyDED-Casp and HyDDCasp (Lasi et al., 2010b) in human HEK-cells. HyDED-Casp has one DED in its N-terminal region and therefore it is similar to mammalian Caspase 8. HyDDCasp has a DD-domain (Lasi et al., 2010a). DD-domains are not typical for animal caspase pro-domains but they occur in some metacaspases of plants and fungi. Expression of HA-tagged versions of these *Hydra* caspases in human HEK-cells revealed a uniform punctate pattern in the cytoplasm for both (Fig. 5B). This pattern changed dramatically, when either caspase was co-expressed with HyFADD-1 or HyFADD-2. Now, the caspases appeared associated with HyFADD-death effector filaments demonstrating that both HyFADD-proteins were able to recruit them (Fig. 6). This suggested that both, HyDED- and HyDD-Caspases can interact with HyFADD-1 or HyFADD-2. Occasionally we observed apoptosis in HEK-cells, when HyDED-caspase was expressed, either alone or together with HyFADD-1 or 2 (Fig. S3 A, C) suggesting that HyDED-caspase, like its human homolog, might also induce apoptosis. However, this effect was not as pronounced as we had previously seen it when human caspase 8 was expressed in HEK-cells (Lasi et al., 2010b) and could not be quantified. Auto-processing of the Hydra DED-caspase with its only DED-domain might not be as efficient in human cells as it is with human caspase 8.

**Fig. 6:**
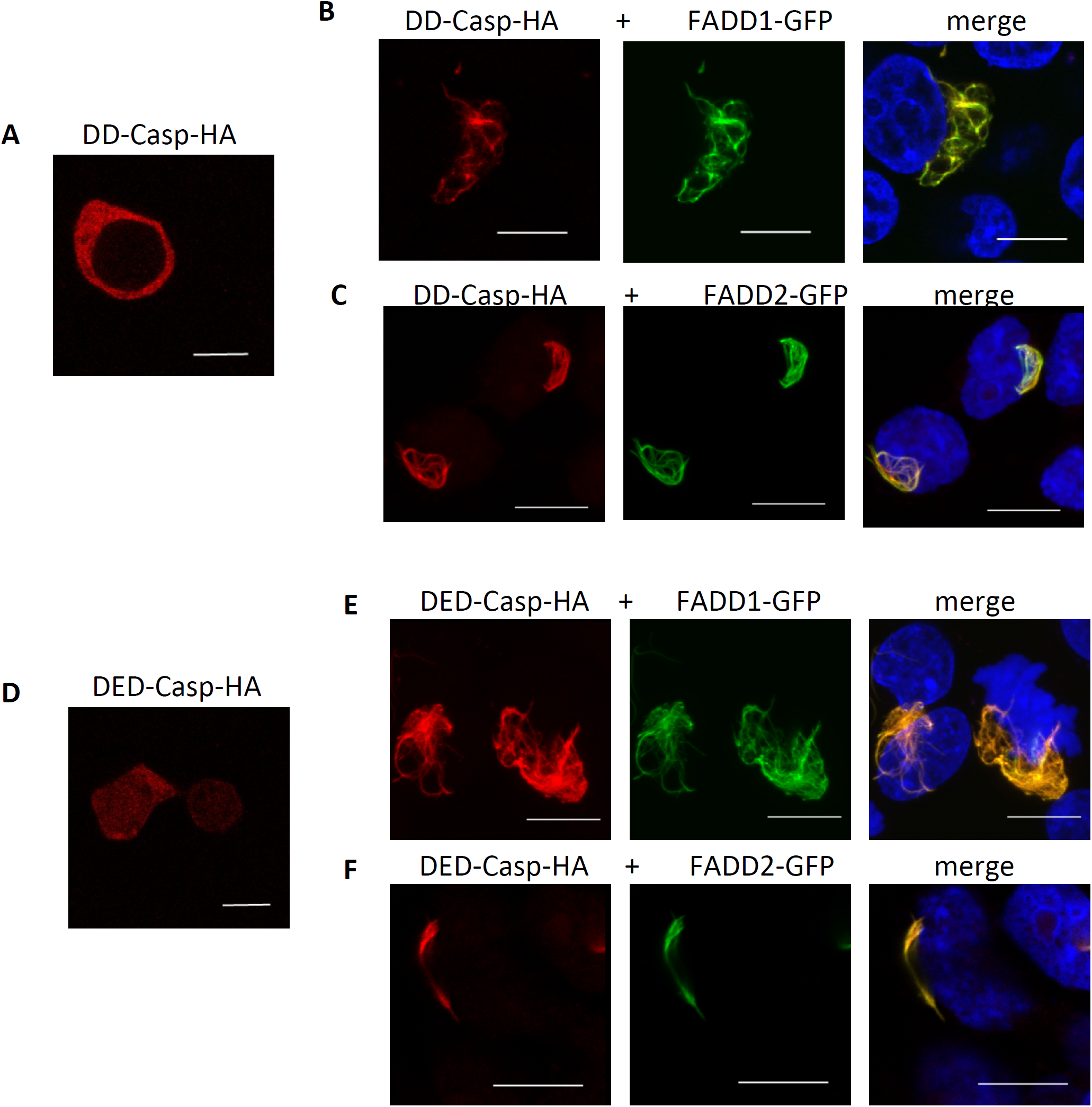
Co-expression of *Hydra* caspases (DD-Casp-HA and DEDCasp-HA) and GFP-tagged HyFADD-1 and HyFADD-2 in human HEK293T cells. Laser confocal images of (A) DD-Casp only (B) DD-Casp co-expressed with HyFADD-1. (C) DD-Casp co-expressed with HyFADD-2. (D) DED-Casp only (E) DED-Casp co-expressed with HyFADD-1. (F) DED-Casp co-expressed with HyFADD-2. Compare A with C and D with F to see complete change of localisation of DED- and DD-casp after co-expression with HyFADDs. GFP (green) anti-HA antibody (red), DAPI (blue); Scale bars: 10 μm.

#### Hydra TNF-receptor does not recruit HyFADD1 and HyFADD2

For control, we co-expressed HyTNF-R with HyFADD-1 and HyFADD-2 in HEK-cells. HyTNF-R was localized in apparent clusters at the plasma membrane and near the nucleus (Fig. 7A). Co-expression of both, HyFADD-1 and HyFADD-2 did not change this distribution of HyTNF-R (Fig. 7B, C). Both HyFADD-proteins were localised in “death effector filaments”, as observed before (Fig. 7B, C) showing no indication of an interaction between HyFADD-proteins and HyTNF-R.

**Fig. 7.**
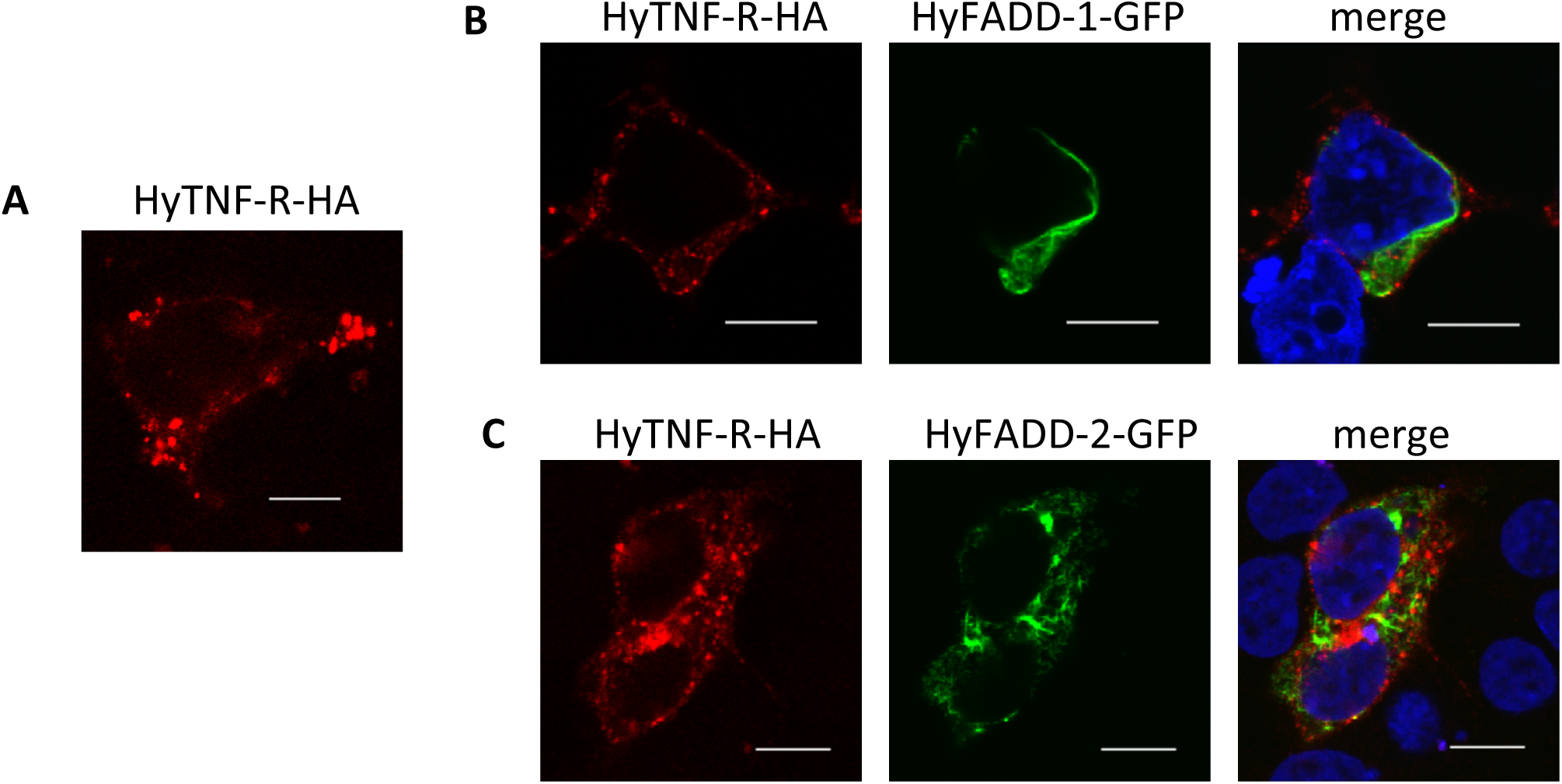
Coexpression of HyTNF-R and HyFADD-1 and −2 in human HEK293T cells. Laser confocal images of (A) HyTNF-R alone. (B) HyTNF-R co-expressed with HyFADD-1. (C) HyTNF-R co-expressed with HyFADD-2 GFP (green) anti-HA antibody (red), DAPI (blue); Scale bars: 10 μm.

**Fig. 8.**
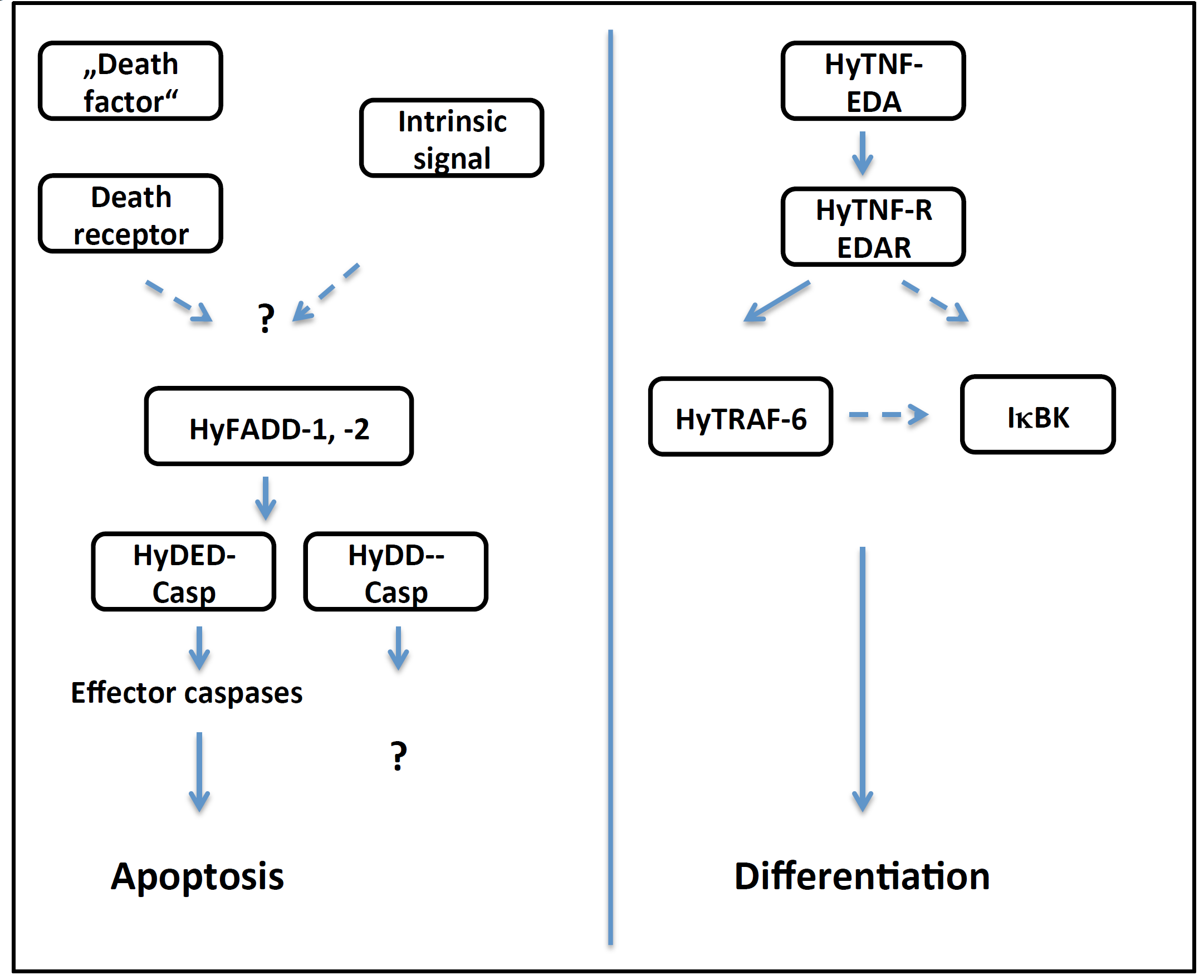
Model for TNF/TNF-R/FADD-function in Hydra: TNF-R (EDAR) mediates differentiation via TRAF-6 and possibly NFκB-signalling; HyFADD1,2 interact with Hy DEDCasp; this might induce apoptosis, the upstream mechanism is unknown. HyFADDs interact with HyDDCasp, upstream and downstream mechanisms are unknown

## Discussion

The induction of intrinsic apoptotic pathways follows a similar molecular logic in vertebrates and invertebrates and this is also true for the pre-bilaterian phylum of cnidaria (Lasi et al., 2010a). In contrast, it is not clear yet, whether the conserved components of extrinsic apoptotic pathways including TNF-Rs, TNFs, DED- and DD-adaptor proteins and Caspase 8, in invertebrates are involved in cell death induction. TNF-R-signalling in mammals is very complex, which is reflected by the large superfamilies of ligands and receptors in this pathway as well as the presence of multiple adaptor molecules that direct signalling at activated receptors into different pathways. Thus, in mammals TNF-R-signalling can either induce or prevent apoptosis and it is involved in many developmental differentiation processes.

As we show here, the hydrozoan *Hydra vulgaris* has only one TNF-R-homolog. Like eight of the *Acropora* TNF-Rs, and in contrast to the single *Drosophila* TNF-R Wengen, HyTNF-R has an intracellular DD-domain. However, the composition of its extracellular domain as well as phylogenetic analyses clearly indicated that it belonged to the EDAR-family of TNF-Rs. We also discovered only one potential TNF-ligand. This has an extracellular TNF-domain followed by a collagen domain, a domain composition as in human EDA - the ligand for EDARs.

In vertebrates, the EDAR-pathway is involved in the development of ectodermal appendages like teeth, hair and glands in mammals, feathers in chicken and scales in fish (reviewed in (Lefebvre and Mikkola, 2014)). The receptor is activated by the TNF-family ligand EDA (reviewed in (Mikkola, 2008)). Ligand binding leads to NFκB activation involving the adaptor molecules EDARADD, TRAF6 and IκB kinase (Sadier et al., 2014). Target genes of this pathway play a role in mediating morphological changes by modulating the actin cytoskeleton and also in the switch from proliferation to growth arrest and differentiation (Kumar et al., 2001)

By immunoprecipitation with anti-TNF-R antibody, we co-precipitated *Hydra* homologs of an IκB kinase subunit and HyTRAF-6. We also co-precipitated *Hydra* homologs of cytoskeletal proteins and thus suggest biochemical interactions of HyTNF-R with components of the mammalian EDAR-signalling cascade. The adaptor protein EDARADD was not found, the gene was also not present in the *Hydra* genome. However, in mammalian 293T and MCF-7 cells it has been shown, that EDAR also interacts with TRAF-6 directly (Kumar et al., 2001). HyTNF-R protein was found very specifically in battery cells and in those epithelial cells of the body column, which have nematocytes incorporated. This suggests that ectodysplasin signalling in *Hydra* functions within in a pathway of epithelial cell differentiation producing nematocyte enclosing cells. Future loss of function studies should be carried out to confirm this.

We did not find any indication of a pro-apoptotic function of HyTNF-R. Moreover, we did not find evidence for an interaction of the HyTNF-R death domain with *Hydra* FADD-homologs, HyFADD1 and HyFADD2. However, expression of HyFADDs in *Hydra* cells seemed to induce cell death. Moreover, when expressed in human HEK293T-cells, HyFADD proteins formed “death effector filaments”, which recruited HyDD-caspase and HyDED-caspase. This suggested that HyDD-caspase as well as HyDED caspase interact with HyFADDs. Occasionally we observed apoptosis in HEK293T-cells, when HyDED-caspase was expressed in HEK-cells, either alone or together with HyFADD-1 or −2 (Fig. S3). Therefore we propose that a FADD-dependent death pathway might exist in *Hydra*. Future experiments will have to reveal how HyFADDs are activated to recruit HyDED- and/or HyDD-caspases and induce apoptosis. This could happen by an extrinsic pathway involving an unknown receptor, but also by intrinsic apoptotic triggers.

We propose that HyTNF-R and HyFADD-proteins are not part of a common extrinsic death pathway. This might be different in the cnidarian *Acropora.* Here a much larger family of TNF-R homologs was found and at least four of the predicted TNF-R proteins have the classical domain structure with transmembrane and intracellular DD-domains as we see it in HyTNF-R. A further two have death domains but no transmembrane domains (Quistad et al., 2014).

Phylogenetic analyses shows that all cnidarian TNF-Rs either group with human EDAR or outside of mammalian TNF-R-, TRAIL- and NGF-R/p75-subfamilies. Of the five *Acropora* TNF homologs two have an extracellular collagen domain, which suggests that they are related to EDA, like HyTNF. This is similar with *Nematostella* TNF-homologs where two of four have a collagen domain ((Robertson et al., 2006) and database search). Thus, in cnidarians we have two subfamilies of the TNF-R superfamily. One is related to mammalian EDARs and, as we show here, probably involved in morphogenesis rather than apoptosis. The function of TNF-Rs outside the EDAR-subfamily in cnidarians should be investigated more closely to confirm that they are involved in extrinsic apoptotic pathways, as it was previously suggested (Quistad et al., 2014).

In the genome of the nematode *Caenorhabditis elegans* neither TNF- nor TNF-receptor encoding genes have been found. *Drosophila melanogaster* does express genes of the extrinsic apoptotic pathway, including the TNF-receptor homolog Wengen and the TNF-homolog Eiger (Kauppila et al., 2003). However, the intracellular domain of Wengen does not have DD-domains. Wengen signalling has been proposed to involve JNK (N-terminal Jun kinase) and to function in host defence, tissue growth, pain sensitization and sleep regulation (reviewed (Igaki and Miura, 2014). In higher invertebrates including ascidians and sea urchins several TNF- and TNF-receptor family members have been identified, which are involved in inflammation and immune response (Parrinello et al., 2008; Romero et al., 2016). This together with our data about *Hydra*-TNF-R provides very clear evidence that non-apoptotic morphogenetic functions are an ancient trait of TNF-R-family members. If and how extrinsic pathways of apoptosis induction work in invertebrates is still an open question.

## Materials and methods

### Gene cloning

Searches for *Hydra* genes with sequence homology to human FADD, TNF, TNF-R, TRAF6 and TRAF4 were performed using the *Hydra* genome server (http://hydrazome.metazome.net/cgi-bin/gbrowse/hydra/) and the NCBI homepage (http://www.ncbi.nlm.nih.gov/). Sequences for HyFADD-1 (XP_002166467.1), HyFADD-2 (XP_002166848.2), HyTNF-R (XP_004209206.1), HyTNF (XP_012554653.1), HyTRAF-6 (XP_004206538.1) and HyTRAF-4 (XP_004207529.1) were amplified from cDNA and cloned into the pSC-B-amp/can vector using the StrataClone Blunt PCR Cloning Kit (Stratagene). For expression in mammalian cells genes were into the vector pHA (clontech), for HA-tagged proteins, and into the vector pEGFP (clontech), for GFP-tagged proteins.

### Cell culture, transfection and immunostaining

Human embryonic kidney (HEK) 293T cells were cultured in Dulbecco’s modified Eagle’s medium (DMEM) supplemented with 10% fetal calf serum, penicillin (100 U ml^−1^) and streptomycin (100 μg ml^−1^) at 37°C, 5% CO_2_. For microscopy, HeLa cells were grown to 50–70% confluence on glass coverslips and transfected with plasmids using Lipofectamine 2000 (Invitrogen) according to the manufacturer’s instructions. 24 hours post-transfection, cells were fixed with 4% paraformaldehyde and permeabilised with 1% Triton-X-100 in phosphate buffered saline (PBS). Anti-HA (H6908, Sigma) was used as primary antibody, Alexa495 anti-rabbit (Invitrogen) was used as secondary antibody.

### *Hydra* culture

*Hydra* were cultured in *Hydra* medium (0.1 mM KCl, 1 mM NaCl, 0.1 mM MgSO_4_, 1 mM Tris-HCl, 1 mM CaCl_2_ (pH 7.6)) at 18 °C. They were fed daily with freshly hatched *Artemia nauplii* larvae and washed after 6-8 h to remove undigested material.

### Generation of antibodies

The peptide (EDIAHFDLSPKTATVDLLHD) injected into rabbits for immunization, the antibody was produced by *Davids Biotechnology*, Regensburg.

### Immunoprecipitation of endogenous HyTNF-R

500 *Hydra* polyps(∼ 5×10^7^ cells) were lysed by pipetting in 500μl solubilization buffer (50 mM Tris, 1% Triton X-100, 1 mM PMSF, 0,1% Bacitracin, 0,2% Aprotinin), left at 4 °C for 30 min and centrifuged at 186 000 g for 1 h at 4 °C. The clear supernatant was incubated at 4 °C for 2 h with 5 μg of anti-HyTNF-R antibody and for control with anti-Hyp53, 50 μl of Protein G Sepharose beads (4 fast flow, GE healthcare) were added for 2 h at 4 °C. After centrifugation the beads were washed four times with wash buffer 1 (150 mM NaCl, 20mM Tris–HCl pH 7.5), four times with wash buffer 2 (300 mM NaCl, 20 mM Tris–HCl pH 7.5) and resuspended for mass spectrometric analysis or PAGE/Western blotting.

### Immunostaining of *Hydra* whole mounts

Polyps were relaxed for 2 min in 2% urethane, fixed with 2% paraformaldehyde for 1 h at room temperature, washed with PBS, permeabilized (0.5% Triton/PBS) and blocked (0.1% Triton, 1% BSA/PBS). Incubation with the primary antibody in blocking solution was carried out overnight at 4 °C. The secondary antibody was incubated for 2 h at room temperature followed by washing and DAPI staining. The animals were mounted on slides with Vectashield mounting medium (Alexis Biochemicals).

### Biolistic transformation

Gold particles (1,0 μm BioRad) were coated with plasmid DNA according to the instructions of the manufacturer. They were introduced into hydra cells with the helios gene gun system (BioRad) as previously described (Böttger et al., 2002)

### Confocal Laser Scanning Microscopy

Leica SP5-2 confocal laser-scanning microscope was used for Light optical serial sections. Leica SP5-2 equipped with an oil immersion Plan-Apochromat 100/1.4 NA objective lens. EGFP and Alexa488 were visualized with an argon laser at excitation wavelength of 488 nm and emission filter at 520 - 540 nm, a UV *laser diode with exitation wavelength of 405 nm and emission filter of* 415 – 465 nm was used for DAPI and Alexa 495 was visualized using a Krypton laser excited at a wavelength of 561nm and emission filter at 604 – 664 nm.

### Domain structure analysis

Domain structures of protein sequences were analysed using the SMART database (http://smart.embl-heidelberg.de/).

### Phylogenetic tree

Neighbour joining trees were calculated using the http://www.phylogeny.fr/ server and Clustal omega https://www.ebi.ac.uk/

## Acknowledgements

This work was funded by DFG-grant BO-1748-7 awarded to AB. We are grateful to Charles David for helpful discussions about this work.

---

**Fig. S1.**
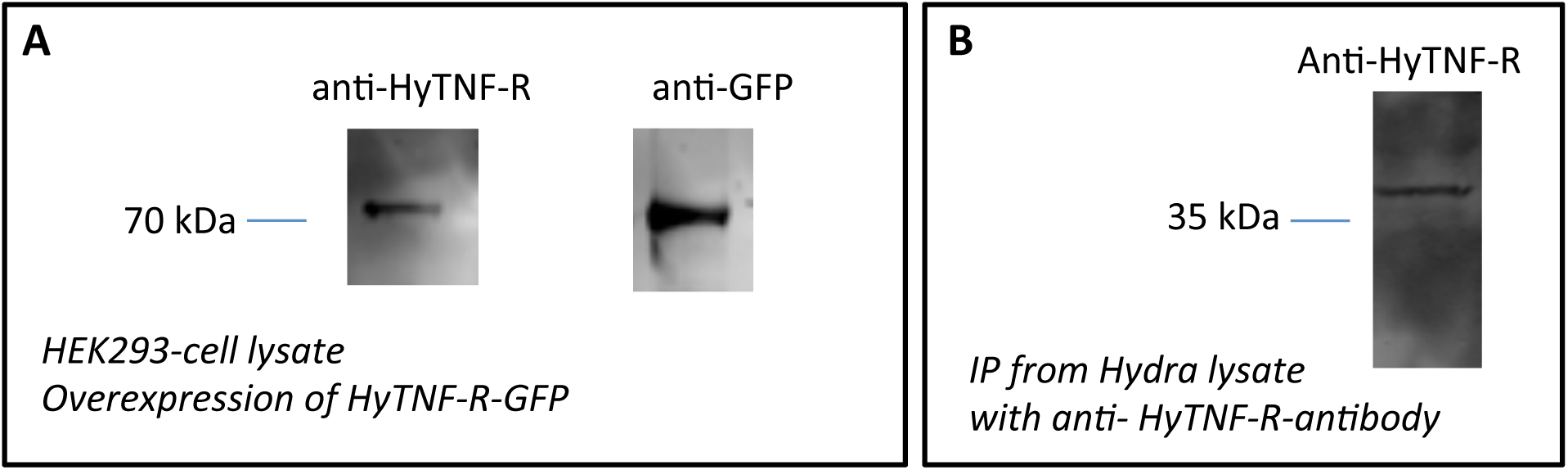
Characterisation of anti-HyTNF-R antibody. (A) Western Blot after PAGE of HEK293T cell lysate expressing GFP-tagged HyTNF-R stained with anti-HyTNF-R and anti-GFP antibodies (B) Western Blot after SDS-PAGE of proteins precipitated with anti-HyTNF-R antibody from *Hydra*-lysate stained with anti-TNF-R antibody

**Fig. S2.**
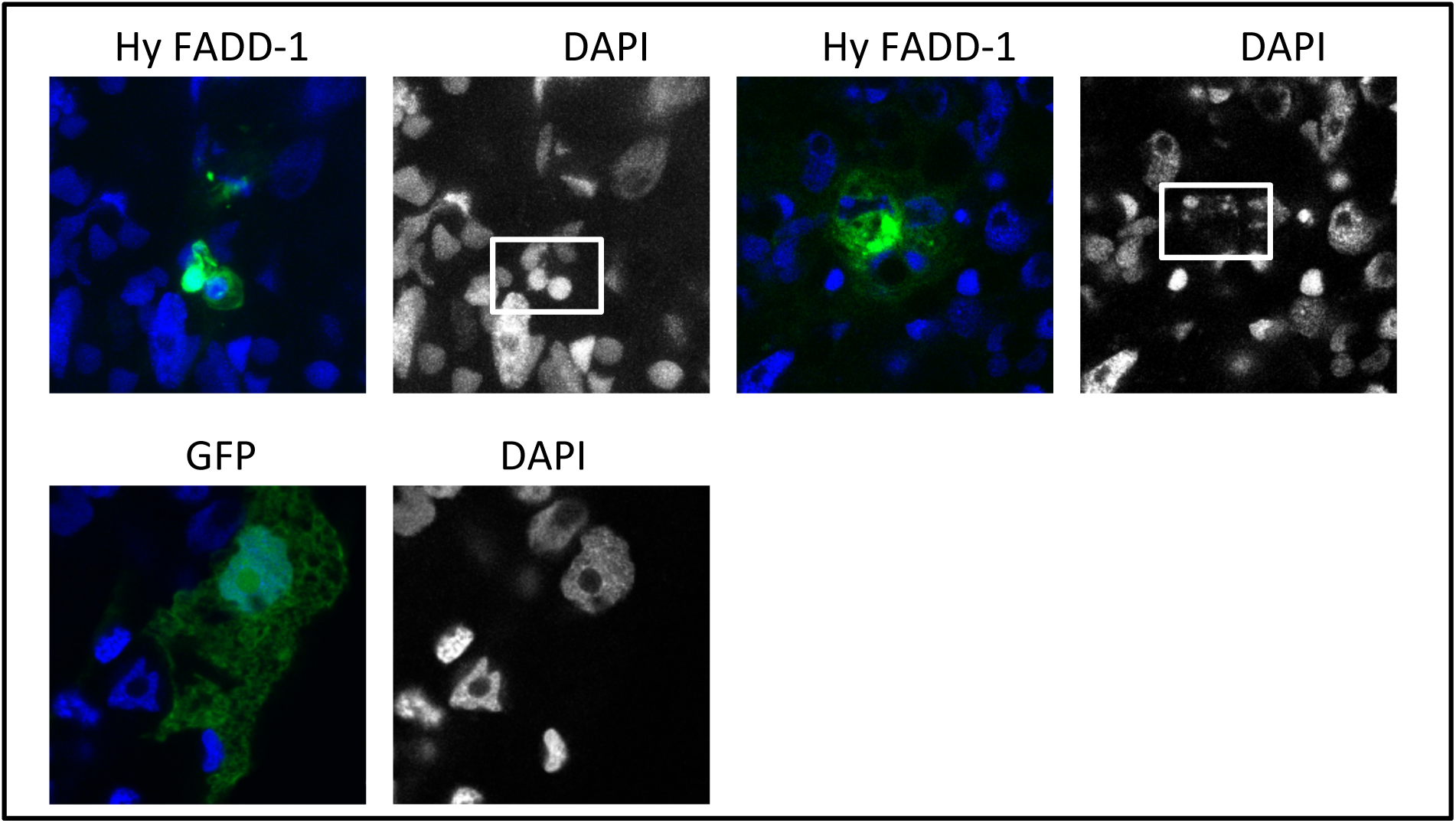
HyFADD1-GFP in *Hydra* cells. Apoptotic cells were observed after expressing HyFADD-1 in *Hydra*, but not after expression of GFP. White boxes indicate apoptotic nuclei. Green: GFP (green), DAPI (blue) in merged image, grey in single image HyFADD1: 2 /2 cells were apoptotic, GFP: 0/100 cells were apoptotic.

**Fig. S3.**
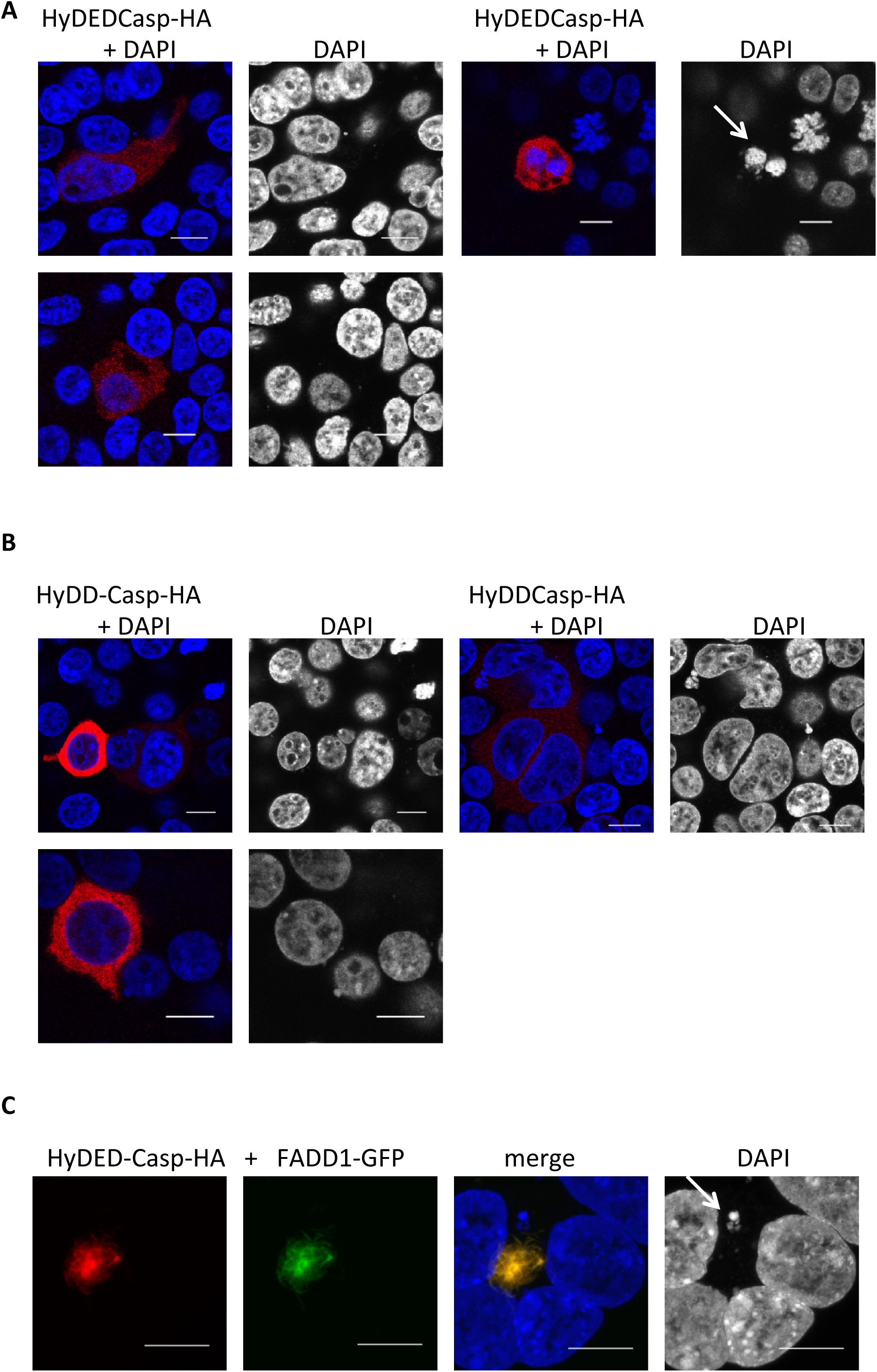
Expression of *Hydra* caspases in HEK293T cells. (A) HyDED-Caspase. (B) HyDD-Caspase. (C) HyDED-caspase co-expressed with GFP-tagged HyFADD-1. White arrow indicates apoptotic nuclei. Green: GFP (green), DAPI (blue) in merged image, grey in single image, HA-tag (red)

**Tab. S1.** List of *Hydra* proteins co-immunoprecipitated with HyTNF-R, probability scores in sample and control-precipitation and manual functional assignments are indicated

| Identified Proteins | Accession Number | Molecular Weight | function | Scaffold probability score |  |
| --- | --- | --- | --- | --- | --- |
|  |  |  |  | sample | control |
| Protein flightless-1 homolog (Fragment) OS=Hydra vulgaris GN=FLII PE=2 SV=1 | T2MD87_HYDVU | 115 kDa | cytoskeleton, actin binding | 100% | 0 |
| Lethal(2) giant larvae protein homolog 1 (Fragment) OS=Hydra vulgaris GN=LLGL1 PE=2 SV=1 | T2MCV2_HYDVU | 165 kDa | cytoskeleton, cell polarity | 100% | 0 |
| Tubulin polymerization-promoting protein family member 2 OS=Hydra vulgaris GN=TPPP2 PE=2 SV=1 | T2MFG9_HYDVU | 19 kDa | cytoskeleton, microtubules | 100% | 0 |
| PPOD1 peroxidase OS=Hydra vulgaris PE=2 SV=1 | Q2FBK4_HYDVU | 32 kDa | ECM | 100% | 0 |
| Pregnancy zone protein (Fragment) OS=Hydra vulgaris GN=PZP PE=2 SV=1 | T2MGN1_HYDVU | 171 kDa | ECM | 100% | 0 |
| Discoidin domain-containing receptor 2 (Fragment) OS=Hydra vulgaris GN=DDR2 PE=2 SV=1 | T2MCH3_HYDVU | 129 kDa | membrane | 100% | 0 |
| Beta-catenin OS=Hydra vulgaris PE=2 SV=1 | Q25100_HYDVU | 90 kDa | membrane | 100% | 0 |
| Nicalin (Fragment) OS=Hydra vulgaris GN=NCLN PE=2 SV=1 | T2M7J4_HYDVU | 63 kDa | membrane | 100% | 0 |
| Splicing factor 3B subunit 3 OS=Hydra vulgaris GN=SF3B3 PE=2 SV=1 | T2MD09_HYDVU | 136 kDa | other | 100% | 0 |
| DNA damage-binding protein 1 OS=Hydra vulgaris GN=DDB1 PE=2 SV=1 | T2MD81_HYDVU | 127 kDa | other | 100% | 0 |
| 116 kDa U5 small nuclear ribonucleoprotein component OS=Hydra vulgaris GN=EFTUD2 PE=2 SV=1 | T2MDI8_HYDVU | 110 kDa | other | 100% | 0 |
| Nuclear cap-binding protein subunit 1 (Fragment) OS=Hydra vulgaris GN=NCBP1 PE=2 SV=1 | T2MEB2_HYDVU | 97 kDa | other | 100% | 0 |
| Uncharacterized protein C21orf63 OS=Hydra vulgaris GN=C21orf63 PE=2 SV=1 | T2MFB8_HYDVU | 43 kDa | other | 100% | 0 |
| Phosphoglucomutase-1 OS=Hydra vulgaris GN=PGM1 PE=2 SV=1 | T2MG15_HYDVU | 60 kDa | other, enzyme | 100% | 0 |
| Nicotinate phosphoribosyltransferase (Fragment) OS=Hydra vulgaris GN=NAPRT1 PE=2 SV=1 | T2MBK0_HYDVU | 70 kDa | other, enzyme | 100% | 0 |
| Phenylalanyl-tRNA synthetase alpha chain (Fragment) OS=Hydra vulgaris GN=FARSA PE=2 SV=1 | T2M2Z6_HYDVU | 58 kDa | other, enzyme | 100% | 0 |
| Protein arginine N-methyltransferase 3 (Fragment) OS=Hydra vulgaris GN=PRMT3 PE=2 SV=1 | T2MHW2_HYDVU | 56 kDa | other, enzyme | 100% | 0 |
| tRNA pseudouridine synthase OS=Hydra vulgaris GN=PUS1 PE=2 SV=1 | T2M305_HYDVU | 50 kDa | other, enzyme | 100% | 0 |
| Adenylosuccinate lyase (Fragment) OS=Hydra vulgaris GN=ADSL PE=2 SV=1 | T2MGT4_HYDVU | 61 kDa | other, enzyme | 100% | 0 |
| NADH-cytochrome b5 reductase OS=Hydra vulgaris GN=CYB5R3 PE=2 SV=1 | T2MHI6_HYDVU | 34 kDa | other, enzyme | 100% | 0 |
| TNF receptor-associated factor 6 (Fragment) OS=Hydra vulgaris GN=TRAF6 PE=2 SV=1 | T2M357_HYDVU | 59 kDa | TNF-R-signalling | 100% | 0 |
| Inhibitor of nuclear factor kappa-B kinase subunit epsilon OS=Hydra vulgaris GN=IKBKE PE=2 SV=1 | T2MDT7_HYDVU | 80 kDa | TNF-R-signalling | 100% | 0 |
| Atlastin-2 OS=Hydra vulgaris GN=ATL2 PE=2 SV=1 | T2M979_HYDVU | 61 kDa | translation/sorting | 100% | 0 |
| Eukaryotic translation initiation factor 5B OS=Hydra vulgaris GN=EIF5B PE=2 SV=1 | T2MHU6_HYDVU | 145 kDa | translation/sorting | 100% | 0 |
| Developmentally-regulated GTP-binding protein 2 (Fragment) OS=Hydra vulgaris GN=DRG2 PE=2 SV=1 | T2MEM0_HYDVU | 52 kDa | translation/sorting | 100% | 0 |
| Probable E3 ubiquitin-protein ligase HERC4 (Fragment) OS=Hydra vulgaris GN=HERC4 PE=2 SV=1 | T2MC32_HYDVU | 57 kDa | translation/sorting endosomes | 100% | 0 |
| Huntingtin-interacting protein 1 (Fragment) OS=Hydra vulgaris GN=HIP1 PE=2 SV=1 | T2MJV4_HYDVU | 39 kDa | translation/sorting, endocytosis | 100% | 0 |
| Protein transport protein Sec24A OS=Hydra vulgaris GN=SEC24A PE=2 SV=1 | T2MH92_HYDVU | 113 kDa | translation/sorting, ER | 100% | 0 |
| Exosome complex exonuclease RRP44 (Fragment) OS=Hydra vulgaris GN=DIS3 PE=2 SV=1 | T2M316_HYDVU | 109 kDa | translation/sorting, exosome | 100% | 0 |
| E3 ubiquitin-protein ligase RNF123 (Fragment) OS=Hydra vulgaris GN=RNF123 PE=2 SV=1 | T2MCA6_HYDVU | 135 kDa | Ub/CARD-domain interaction | 100% | 0 |

